# Effect of cysteine oxidation in SARS-CoV-2 Spike protein on its conformational changes: insights from atomistic simulations

**DOI:** 10.1101/2024.10.24.620034

**Authors:** Maryam Ghasemitarei, Hoda Taeb, Tayebeh Ghorbi, Maksudbek Yusupov, Tapio Ala-Nissila, Annemie Bogaerts

## Abstract

This study investigates the effect of cysteine (Cys) oxidation on the conformational changes of the SARS-CoV-2 Spike (S) protein, a critical factor in viral attachment and entry into host cells. Using targeted molecular dynamics (TMD) simulations, we explore the conformational transitions between the down (inaccessible) and up (accessible) states of the SARS-CoV-2 S protein in both its native and oxidized forms. Our findings reveal that oxidation significantly increases the energy barrier for these transitions, as indicated by the work required to move from the down to the up conformation and vice versa. Specifically, in the oxidized system compared to the native system, the energy required to transition from the down to the up conformation increases by approximately 131 ± 1 kJ.mol^−1^, while the energy required for the reverse transition increases by about 223 ± 6 kJ.mol^−1^. This is due to the stabilizing effect of oxidation on the conformation of the SARS-CoV-2 S protein.

Analysis of hydrogen bond and salt bridge formation before and after oxidation provides additional insights into the stabilization mechanisms, showing an increase in salt bridge formation that contributes to conformational stabilization. These results underscore the potential of targeting translational modifications to hamper viral entry or enhance susceptibility to neutralization, offering a novel perspective for antiviral strategy development against SARS-CoV-2.

This study adds important knowledge to the field of viral protein dynamics and highlights the critical role of structural and computational biology in uncovering new therapeutic avenues.

## Introduction

The emergence of the novel coronavirus, SARS-CoV-2, in late 2019 marked the onset of a global health crisis. Initially identified in Wuhan, China, this virus rapidly spread worldwide, causing the coronavirus pandemic 2019 (COVID-19). Characterized by symptoms such as fever, dyspnea, dry cough, and tiredness, SARS-CoV-2 has affected millions of people [1], resulting in significant illness and mortality. As a member of the coronavirus family, SARS-CoV-2 is the seventh coronavirus known to infect humans and belongs to the β-coronavirus group [2]. Unlike SARS-CoV and MERS-CoV, which have higher mortality rates, SARS-CoV-2 poses unique challenges due to its rapid transmission [3], resulting in a global pandemic.

Coronaviruses are defined by two groups of proteins: structural and non-structural [4]. Among the structural proteins, the Spike (S) glycoprotein plays a crucial role in virus attachment and entry into host cells [5]. Due to its surface exposure and central role in viral pathogenesis, the S glycoprotein is a major target for therapeutic interventions and vaccine development [6]. However, the extensive coverage of S glycoproteins with N- and O-linked glycans presents challenges in targeting specific epitopes for neutralizing antibodies [7, 8].

The SARS-CoV-2 S glycoprotein is a trimeric protein, with each monomer containing two functional subunits: S1 and S2. The S1 subunit is responsible for binding to the host cell receptor, while the S2 subunit facilitates the fusion of the viral and cellular membranes [9]. The S1 subunit comprises four domains: the N-terminal domain (NTD), receptor-binding domain (RBD), and two C-terminal domains (CTD1 and CTD2). The RBD, located in the distal domain of the S1 subunit, helps stabilize the prefusion state of the S2 subunit, which contains the fusion machinery [10]. To interact with a host cell receptor, the RBD of the S1 subunit undergoes hinge-like movements, transitioning between “down” (inaccessible) and “up” (accessible) conformations (see Figure 1) [11, 12]. These conformational changes are crucial for the engagement of host cell receptors [12] by the SARS-CoV-2, SARS-CoV, and MERS-CoV viruses [10].

**Figure 1.**
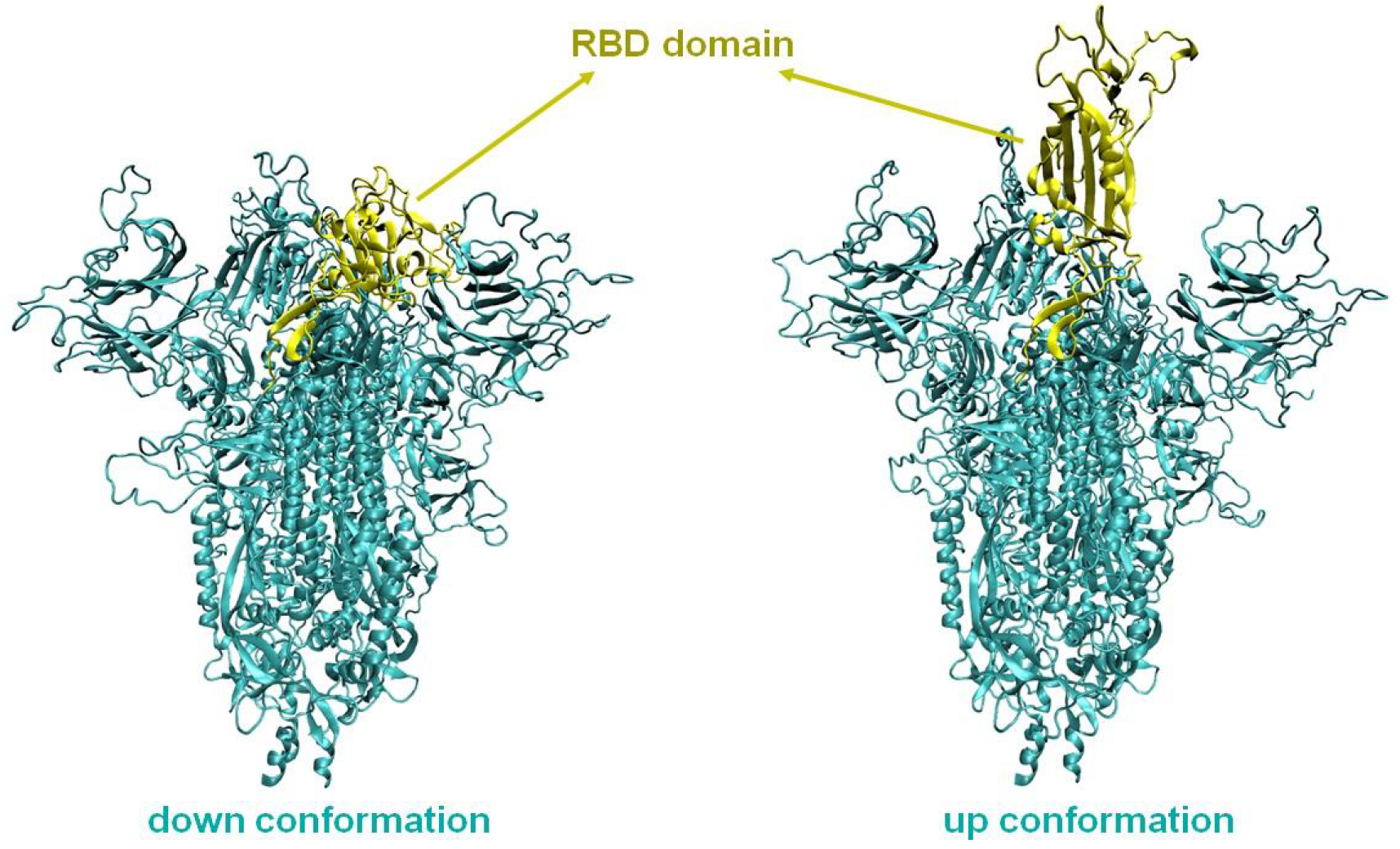
SARS-CoV-2 S protein in the down state (left) and up state (right). The RBD domain is depicted in yellow, while the rest of the SARS-CoV-2 S trimer is shown in cyan.

Each unit of the SARS-CoV-2 S protein contains 19 N-linked and 1 O-linked glycosylation sites, i.e., reacting sites with glycans [13, 14]. Molecular dynamics (MD) simulations have underscored the importance of these glycans in inducing conformational changes in the SARS-CoV-2 S protein [8, 15-17]. The conformational masking and glycan shielding of the SARS-CoV-2 S glycoprotein in the down conformation reduce its recognition by the immune response of the infected host [12, 15].

Generally, two classes of RBD exposure have been reported for the wild type of the S trimer, i.e., all RBDs down, or one up [18]. A 1:1 ratio of these two classes has been observed using cryo-EM [19]. During the transition from the down to the up conformation, the fusion-peptide proximal region (FPPR) clashes with the CTD1, which rotates outward along with the RBD. Therefore, a structured FPPR plays an important role in stabilizing the down conformation [20], although it occasionally flips out of position, allowing the RBDs to move into the up conformation.

It would be highly favorable to stabilize the SARS-CoV-2 S trimer in the down conformation, to impede viral attachment and entry [18, 20-22] in the prefusion state. Studying the details of the SARS-Cov-2 S structure in the down conformation could aid in devising effective methods to keep the SARS-CoV-2 S trimer in this state. In the down conformation, the RBD is oriented closer to the trimer’s central cavity [11], facilitated by strong electrostatic interactions between negatively charged residues like aspartate 427 and 428 (Asp_427_ and Asp_428_), and positively charged lysine 986 (Lys_986_) [18]. Alterations in key residues through mutation can modify the conformational distribution of the S domains, stabilizing the SARS-CoV-2 S trimer in its down structure [19]. For instance, mutations of serine 383 (Ser_383_), Asp_985_, glycine 413 (Gly_413_), and valine 987 (Val_987_) to cysteine (Cys) form disulfide bridges that prevent conformational changes and RBD exposure [18, 19]. Additionally, replacement of Lys_986_, Val_987_ [11], phenylalanine 817 (Phe_817_), alanine 892, 899 and 942 (Ala_892_, Ala_899_ and Ala_942_) with proline (Pro) further stabilizes the SARS-CoV-2 S trimer in its down conformation [23]. Furthermore, mutations of Asp_614_ to asparagine (Asn) and arginine 682 and 685 (Arg_682_ and Arg_685_) to Ser aim to enhance attractive interactions between the RBD and the rest of the SARS-CoV-2 S trimer to maintain the down form [24].

Moreover, certain small molecules like formaldehyde [25], linoleic acid [26-28], and an ultra-potent synthetic nanobody Nb6 [29] can stabilize the SARS-CoV-2 S trimer in its down conformation by creating specific cross-links between RBD protomers and S2 subunit residues. Developing methods to trap the SARS-CoV-2 S trimer in its down state could assist in treating SARS-CoV-2, by preventing RBD attachment to cell receptors and subsequent viral entry. Cold atmospheric plasma (CAP) could also be beneficial in this regard, by oxidizing exposed domains of the SARS-CoV-2 S protein, as part of the mentioned strategy [30].

Over the past decade, CAP has been used to treat various biological systems, including proteins [31], cell membranes [32], and nucleotides [33], resulting in structural oxidation and nitration. CAP treatment of COVID-19 can alter the structure of the SARS-CoV-2 S protein [34], affecting the conformational transition of the RBD. The chemical modification of amino acids by CAP in aqueous solution has been studied in previous research [35, 36]. After CAP treatment, chemical modifications were observed in 14 out of 20 amino acids using high-resolution mass spectrometry [35]. The oxidation of the electron-rich groups in the side chains of amino acids due to CAP-produced reactive species can generate various types of products, including: (i) sulfoxidation of methionine (Met), (ii) cystine formation and sulfonation of thiol groups in Cys, (iii) hydroxylation and nitration of aromatic rings in tyrosine (Tyr), Phe and tryptophan (Trp), (iv) amidation and ring opening in histidine (His) and Pro, and (v) no oxidation in six amino acids, namely Gly, Ser, threonine (Thr), Asn, and Asp [35]. Previous studies have shown that after sulfur-containing amino acids like Met and Cys, aromatic amino acids such as Tyr, Phe and Trp are the next targets for oxidation by reactive species produced by CAP [37].

Investigating the conformational transition of the SARS-CoV-2 S trimer at the atomic scale helps identify key residues that significantly influence these conformational changes. Mutation and/or oxidation of these residues as a result of CAP treatment may interfere with the normal exposure of the SARS-CoV-2 RBD, rendering it inaccessible to the cell receptor and ultimately reducing cell infection [34].

In this study, we investigate the conformational changes of the SARS-CoV-2 RBD from the down to the up conformation using targeted molecular dynamics (TMD) simulations, identifying important residues involved in these transitions. By altering specific Cys residues, which are easily oxidized during CAP treatment [35, 36, 38, 39], we examine the potential disruption of normal conformation of the SARS-CoV-2 RBD and its accessibility to the receptor. Furthermore, the most significant amino acid residues and glycans playing an important role in conformational changes in both the native and oxidized states are identified and compared.

### Computational details

The SARS-CoV-2 S trimer is a large molecule, with each monomer (chains A, B, and C) comprising 1273 amino acid residues. Different conformations of its down and up states are available in the Protein Data Bank (PDB). We selected the cryo-EM structures of the SARS-CoV-2 S trimer for the down and up conformations, corresponding to PDB IDs 6VXX [12] and 6VSB [11], respectively. Since the RBD part has numerous missing residues in both PDB structures, we used the crystal structure of the RBD from PDB ID 6M0J [40] to remodel the structures.

To glycosylate the SARS-CoV-2 S trimer, we employed the CHARMM-GUI web server [41-43]. Based on the literature [12, 13], we selected the most probable glycan structures for both systems. Details of the selected glycans can be found in Table S1 of the Supporting Information (SI).

After preparing the native down and up conformations of the SARS-CoV-2 S trimer, MD simulations were conducted using the Gromacs 2020.2-MODIFIED software [44] to equilibrate both structures. The CHARMM36 force field [45, 46] was employed to describe the interatomic interactions, and the initial structures (model systems) were prepared using the CHARMM-GUI web server [41].

The simulation box dimensions were 201 × 192 × 198 Å^3^ for the down system and 192 × 185 × 204 Å^3^ for the up system. Periodic boundary conditions were applied in all three spatial directions. The charge of both systems was neutralized by adding chloride and sodium ions to the surrounding water, and explicit water molecules were simulated using the TIP3P model [47]. Energy minimization was carried out using the steepest descent algorithm for 20000 steps.

For both systems (i.e., native down and up conformations), three replicas were prepared with different initial atomic velocities using different random seeds. Equilibration was then performed in the NVT ensemble at 310 K for 3 ns with a time step of 1 fs. Subsequently, they were relaxed in the NPT ensemble at 1 atm and 310 K for 200 ns with a 2 fs time step (see the root mean square deviation (RMSD) plots in Figure S1A of the SI). The Nosé-Hoover thermostat [48-50], with a coupling constant of 1 ps and the isotropic Parrinello-Rahman barostat [51], with a coupling constant of 5 ps and a compressibility of 4.5×10^−5^ bar^−1^, were used to equilibrate the systems.

To maintain the orientation of the SARS-CoV-2 S trimer within the simulation box, the tails of the SARS-CoV-2 S trimer were restrained using a harmonic potential with a force constant of k = 4000 kJ.mol^−1^.nm^−2^, specifically restraining the heavy atoms in residues 1140-1147. Long-range interactions were computed using the Particle Mesh Ewald (PME) method [52, 53] with long-range dispersion corrections applied for both pressure and energy. The Verlet list scheme was employed, with a 12 Å cutoff for both electrostatic and van der Waals (VDW) interactions. Considering the cutoff value, interactions occurring within a 12 Å radius between the RBD domain of chain A and the surrounding amino acid residues in the down conformation of the SARS-CoV-2 S trimer were assumed to influence structural changes (highlighted in red in Figures 2A and 2B). During CAP treatment, modifications to amino acid residues within this domain could potentially impact conformational changes. Moreover, the conformational masking and glycan shielding observed in the down conformation of the SARS-CoV-2 S trimer may prevent amino acids from being exposed to solvent and hence reactive oxygen species (ROS) generated by CAP. Thus, to identify the amino acid residues most susceptible to oxidation within this domain, a solvent accessible surface area (SASA) analysis was conducted using the *gmx sasa* tool in Gromacs on the last 10 ns of all three replicas of the native down conformations. The SASA analysis was focused on identifying amino acids within the domain that are highly accessible to the solvent, as these are more likely to undergo oxidation during CAP treatment. However, as there are no Met residues in the selected domain, Cys residues, which can be oxidized to cysteic acid (CYO), were considered next in susceptibility to oxidation. Based on the SASA results (Figure 2C), Cys_480_ in the RBD domain, Cys_166_ in chain B, and Cys_480_ and Cys_488_ in chain C were identified as susceptible to be modified to CYO.

**Figure 2.**
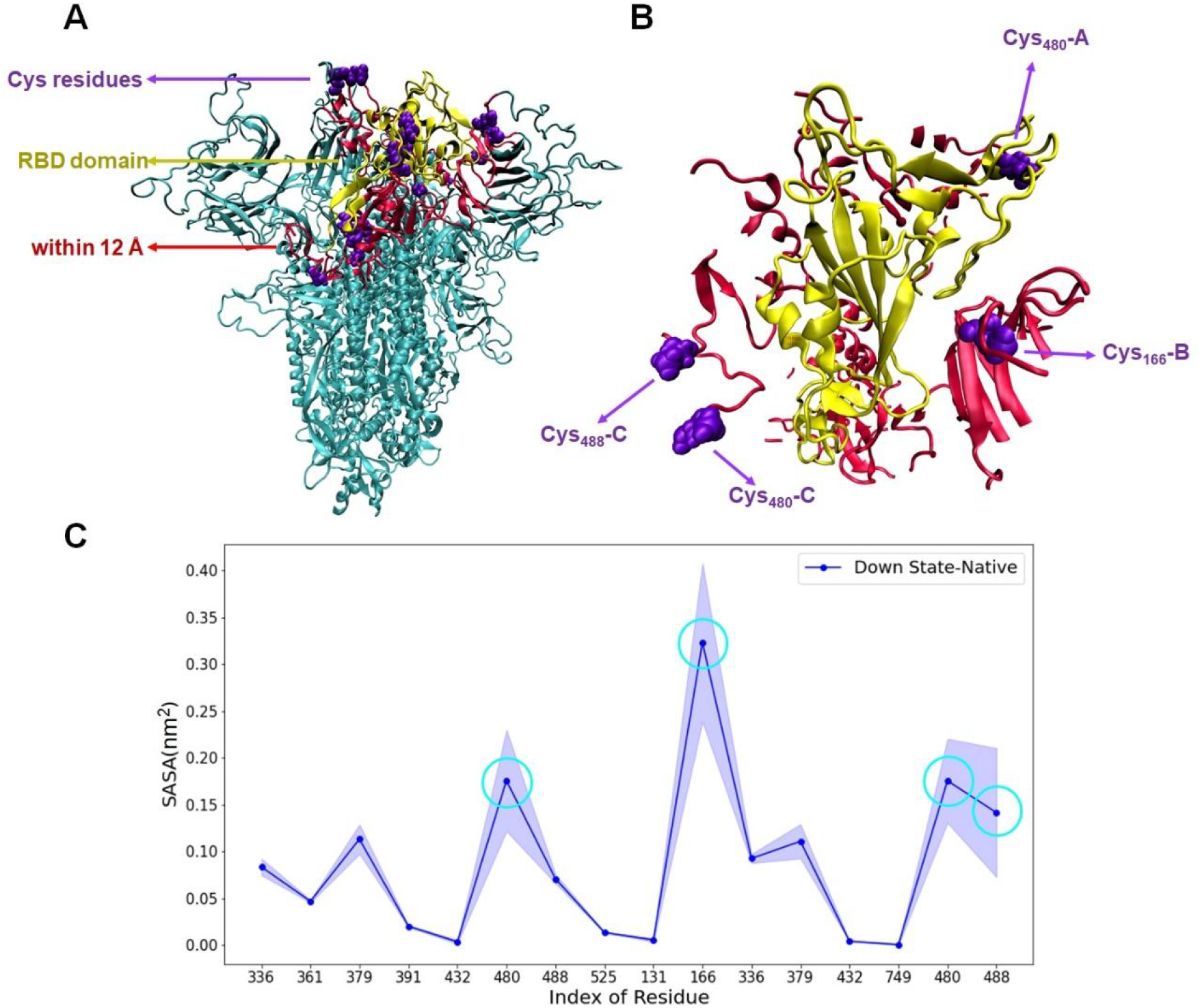
Structural and solvent accessibility analysis of the SARS-CoV-2 S protein in the down state. (A) The RBD domain is highlighted in yellow, with its immediate surrounding region within a 12 Å radius shown in red to emphasize its structural context. All cysteine residues within these domains are marked in purple and represented with VDW spheres. (B) Detailed view of the RBD and its surrounding region within a 12 Å radius, focusing on the areas highlighted in yellow and red. Four cysteine residues demonstrating the highest solvent accessibility, determined by SASA analysis, are again depicted in purple with VDW representation. (C) SASA analysis plot illustrating the solvent accessibility of all cysteine residues within the yellow and red domains. The four cysteines with the highest SASA values are highlighted with light blue circles, indicating their potential importance and susceptibility to oxidation based on detailed analysis.

The partial charges of CYO residues were identified by determining their protonation states in a physiological environment. The negative partial charge of CYO under physiological conditions is given by the DrugBank database [54]. Additionally, experimental investigations on Cys oxidation have also confirmed that the oxidation of Cys residues in proteins, induced by ROS, predominantly results in negatively charged CYO residues [55].

For negatively charged CYO force-field parameters, a combination of Gaussian 16 software [56] and the CHARMM General Force Field (CGenFF) [57] was employed, following [58]. Density Functional Theory (DFT) was applied to optimize and generate the partial charges for CYO using the B3LYP functional with the standard 6-311G* basis set in Gaussian. The resulting partial charges and optimized structure were used to construct topology files for CYO, compatible with the CHARMM36 force field, using the CGenFF program.

Similar to the methods used for the native systems, both the down and up conformations of the oxidized SARS-CoV-2 S trimer were constructed and equilibrated (see the RMSD plots in Figure S1B of the SI). Thus, three replicas were generated for each of the following systems: (i) native down S trimer, (ii) native up S trimer, (iii) oxidized down S trimer, and (iv) oxidized up S trimer. This resulted in twelve systems in total.

Using PLUMED-2.6.0 software [59-61], TMD simulations were performed to transition the SARS-CoV-2 S trimer between its down and up conformations and vice versa. From the last 100 ns of trajectories, three structures from all three replicas were selected as an initial and target conformations for the TMD simulations, resulting in a total of nine structures for each system (i.e., native down, native up, oxidized down, and oxidized up S trimers). During the TMD simulations, each of the nine systems in the down state was randomly targeted to one of the nine structures in the up state, and vice versa. Consequently, for each native and oxidized system, nine TMD simulations were conducted for the transition from down to up, and another nine TMD simulations were performed for the transition from up to down. Each TMD simulation was run for 5 ns to guide the SARS-CoV-2 S trimer from the down to the up state. This was achieved by applying a spring with a constant free length *ρ* = 0.05 Å and a linearly increasing elastic constant ranging from 0 to 80000 kJ.mol^−1^. nm^−2^. In the TMD simulations, all heavy atoms were subjected to spring forces to guide them from the initial to the final structure. The work of transition during the TMD simulation is defined by:

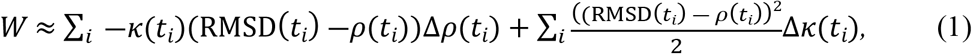

where Δ*ρ*(*t*_*i*_) and Δ *κ*(*t*_*i*_) denote the changes in the center of the harmonic potential and the spring constant in the *i*th step, respectively.

To investigate the pathway between up and down states, the VMD software [62] was employed. Initially, the number of hydrogen bonds (H-bonds) and salt bridges between the RBD and its surroundings in the down conformation of the SARS-CoV-2 S trimer were computed. Subsequently, differences in the pathways of H-bond and salt bridge dynamics during the down-to-up transition between the native and oxidized systems were compared. Quantitative analyses of H-bonds and salt bridges were conducted based on the methodology described by Debiec et al. [63], adopting specific criteria: a distance of 3.5 Å for salt bridge formation and 4.5 Å for dissociation, along with a 3 Å cutoff distance for hydrogen bond formation [64]. The quantities of H-bonds and salt bridges were determined by averaging their occurrence over the last 100 ns of MD simulations, where each interaction was prevalent in more than 10% of all frames.

## Result and discussion

The total work of the applied force needed for transition between two states is calculated for both native and oxidized systems. The work difference, Δ W = W_*oxidized*_ - W_native_, is presented in Figures 3A and 3B. The results indicate that the work required for the transition from the down to the up conformation and vice versa was higher in the oxidized system as compared to the native system. Specifically, the difference in work for the transition from the down to the up state is 131 ± 1 kJ.mol^−1^, and from the up to the down state it is 223 ± 6 kJ.mol^−1^. These findings suggest that Cys oxidation makes the transition barriers higher (both in down to up and up to down) and makes the transition less probable (more difficult) in the oxidized system as compared to the native one, resulting in increased stability of the SARS-CoV-2 S protein in each state.

**Figure 3.**
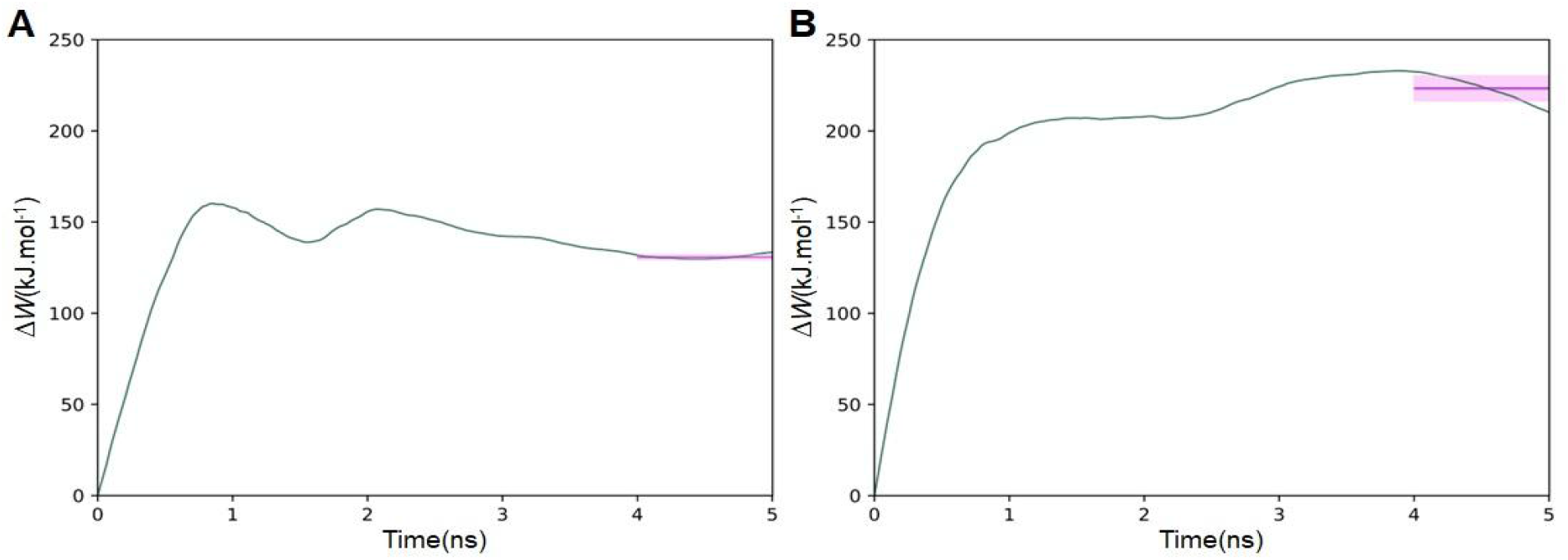
The difference in the work associated with the TMD applied force between the oxidized and native systems during down-to-up transition (A), and up-to-down transition (B). The purple curve illustrates the averaged value for the last one nanosecond of the TMD simulations.

The RMSD values of the RBD during the transition from down to up state revealed that in the oxidized system there are different regimes of transitions (Figure S2 in the SI). The averaged RMSD slope distinguishes between rapid and slow transition regimes. For the first 1.5 ns, the system undergoes rapid changes, while from 1.5 to 4 ns, it moves much slower toward the final state. A detailed investigation of the RMSD is needed to realize the behavior. As stated in Ref. [34], the RBD can be indicated in 5 non-overlapping regions: I (Cys_336_–Cys_361_), II* (Cys_379_– Leu_390_), II&III (Cys_391_–Cys_432_), III* (Val_433_–Pro_479_ and Tyr_489_–Cys_525_), and IV (Cys_480_–Cys_488_).

Regions III* and IV are particularly crucial as they interact with host cell receptors for cell entry and fusion. The RMSD of these 5 regions during the transition for both the native and oxidized systems show that the two distinct phases observed in the oxidized system are associated with these two domains. Figures 4A and 4B illustrate the RMSD values of these 5 regions during the transition from the down to the up state for the native and oxidized system, respectively. The magenta and green curves represent regions III* and IV, respectively, which exhibit significant changes in the oxidized system compared to the native one, highlighting their role in the observed two-regime of RMSD behavior in the oxidized system.

**Figure 4.**
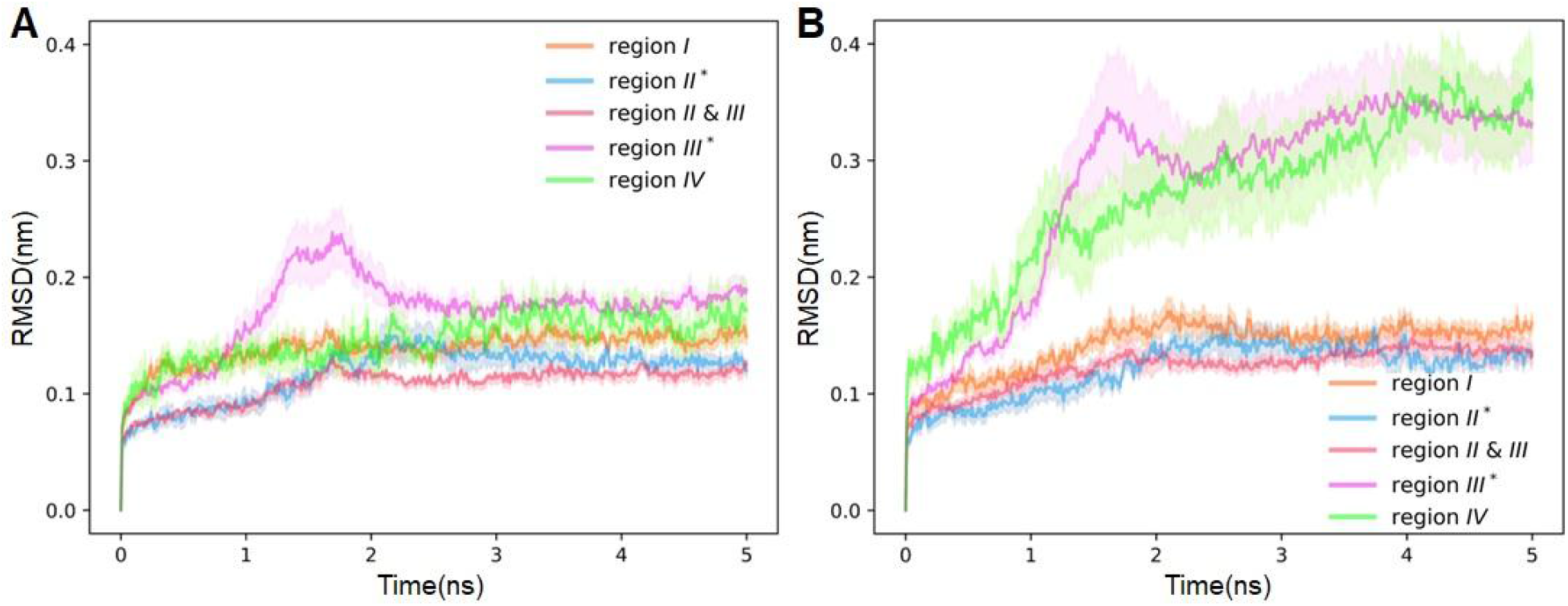
RMSD of each region of the native (A) and oxidized (B) RBD of the SARS-CoV-2 S trimer during down-to-up transition, all averaged over nine replicas.

Further investigation into the trajectories of TMD simulations revealed that glycans, notably N165B, N343A, and N343B, play a critical role in the slower transition regime with higher work values observed in the oxidized RBD. Analysis of H-bonds in the down state of both systems showed that in the native system, five glycans, N331A, N343A, N165B, N234B, and N343B, significantly contribute to forming H-bonds (Tables S2, S3, and S4 in the SI), whereas in the oxidized system, only N234B and N343B form H-bonds with the RBD. During the transition, the presence of negatively charged CYO residues plays a key role in attractive interactions with these glycans. In both systems, N234B and N331A, located at the hinge domain, remained stable relative to RBD movements. However, N165B was pulled upwards by the RBD in both the native and oxidized systems. A notable difference between the two systems was observed with N165B, which detached from the RBD around 1.6 ns in the TMD simulation of the native system but remained attached throughout the entire TMD simulation in the oxidized system. Figures 5A and 5B depict the positions of these glycans and the RBD domain during the transition from the down to the up state in the native and oxidized systems, respectively. From these figures, it is evident that there were no significant differences between the two systems in terms of N234B and N331A (depicted in white and licorice representations). The distinction primarily lies in the interactions involving oxidized Cys residues (depicted in VDW representation) and glycans N343A and N343B. Specifically, N343A in the RBD was attracted to CYO_480_ and CYO_488_ of chain C, hindering easy movement of the RBD. Moreover, CYO_480_ in the RBD, initially not attracted to any glycans, found N343B during the transition and rotated towards it, preventing complete upward movement. The second phase of RMSD observed after approximately 1.5 ns could be attributed to these interactions, as CYO_480_ is positioned at the boundary between region III* and region IV. To visualize the dynamic behavior of the system, trajectories from TMD simulations were converted into video format.

**Figure 5.**
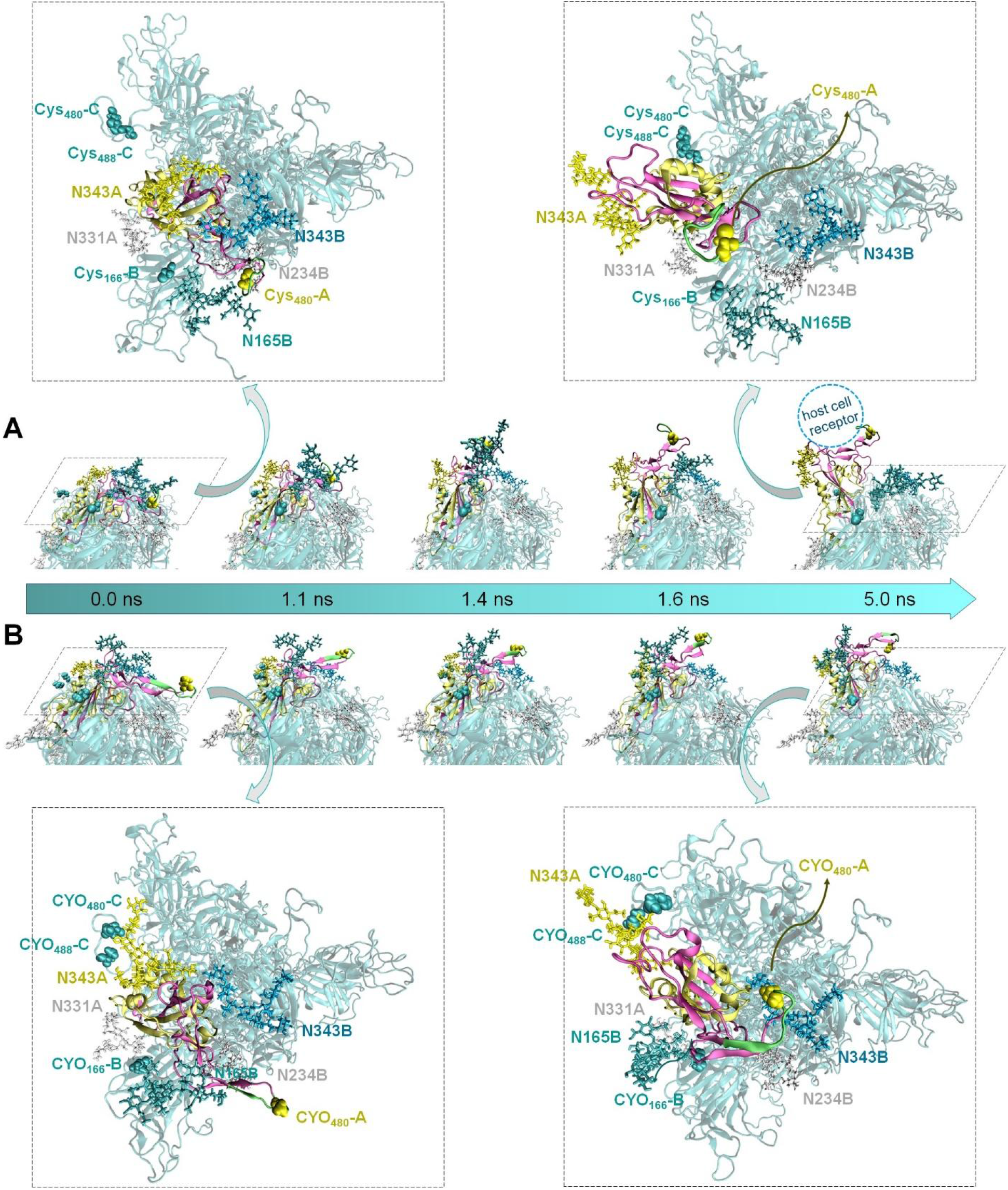
The position of RBD decorating glycans in relation to the rest of the SARS-CoV-2 S trimer during down-to-up transition for the native (A) and oxidized (B) systems. Region III*, IV, and the rest of the RBD are illustrated in magenta, green, and yellow colors, respectively. The rest of the SARS-CoV-2 S trimer, is depicted in cyan color. The glycans N331A and N234B are represented in white using licorice representation. The concerned Cys residues in the oxidized system (see text) are shown with VDW representation. To enhance clarity on the positions of the RBD, glycans, and (native/oxidized) concerned Cys residues, the top views of both the initial (0 ns) and final (5 ns) states of each system are zoomed in. The position of the host cell receptor is indicated by a dashed circle. It is evident from the final state of the oxidized system that the RBD is less exposed for host cell receptor attachment compared to the native form.

These videos, available online at https://github.com/maryamghasemitarei/Transion-of-SARS-CoV-2-Spike-protein, offer insights into the conformational changes and interactions, helping in a deeper understanding of the structural evolution during the transition.

The average total number of H-bonds and salt bridges formed between the RBD domain and the remaining SARS-CoV-2 S trimer (with a prevalence exceeding 10% in all frames) is summarized in Tables 1 and 2, respectively. The results indicate that after oxidation, there was no significant change in the number of H-bonds, but there was an increase in the number of salt bridges. This increase suggests that the transition process becomes more challenging in the oxidized protein.

**Table 1.** Number of H bonds formed between RBD and each chain of SARS-CoV-2 S protein (chains A, B and C), along with the total H bonds. Values are averaged over the last 100 ns of MD simulation, representing bonds present in more than 10% of the frames.

|  |  | Native | Oxidized |
| --- | --- | --- | --- |
| RBD | chain A | $5.42 \pm 0.26$ | $5.43 \pm 0.24$ |
| | chain B | $4.64 \pm 0.30$ | $5.12 \pm 0.46$ |
| | chain C | $0.79 \pm 0.11$ | $0.64 \pm 0.12$ |
| | total | $10.85 \pm 0.41$ | $11.19 \pm 0.53$ |

**Table 2.** Number of salt bridges formed between RBD and each chain of SARS-CoV-2 S protein (chain A, B and C), along with the total number of salt bridges. Values are averaged over the last 100 ns of MD simulation, representing bridges present in more than 10% of the frames.

|  |  | Native | Oxidized |
| --- | --- | --- | --- |
| RBD | chain A | $1.40 \pm 0.19$ | $1.95 \pm 0.02$ |
| | chain B | $0.70 \pm 0.21$ | $1.11 \pm 0.13$ |
| | chain C | $0.18 \pm 0.13$ | $0.12 \pm 0.06$ |
| | total | $2.28 \pm 0.31$ | $3.18 \pm 0.14$ |

In the interface between all pairs forming H-bonds (Tables S2, S3, and S4 in the SI), a total of fourteen pairs were identified within the hinge domain. These pairs exhibited minimal movement during the transition from the down to the up conformation, and the distance between their donor and acceptor atoms did not significantly change during this transition (Figure S3 in the SI). During the transition from the down to the up state, the remaining H-bonds between RBD and its surrounding SARS-CoV-2 S trimer were broken and the distance between donor and acceptor atoms in both the native and oxidized systems were increased. Notably, the oxidized system exhibited a slower progression, indicated by a reduced slope in the distance versus time plot compared to the native system (Figure 6). Detailed distance measurements for three representative donor-acceptor pairs are shown in Figure 6C, where for additional pairs, they are presented in Figure S3 in the SI. Similarly, among all pairs forming salt bridges (Table S5 in the SI), four pairs – Arg_328_-Asp_578_, Arg_319_-Asp_737_, Lys_535_-Glu_583_, and Glu_471_-Lys_113_ – were identified within the hinge domain, with minimal change in nitrogen-oxygen distance during the transition (Figure S4 in the SI). However, salt bridges in other pairs – Lys_462_-Asp_198_, Lys_535_-Glu_554_, and Asp_427_-Lys_986_ – were disrupted during the transition. Consistent with the H-bond results, comparison of the distance versus time slopes indicated a slower transition in the oxidized system compared to the native system.

**Figure 6.**
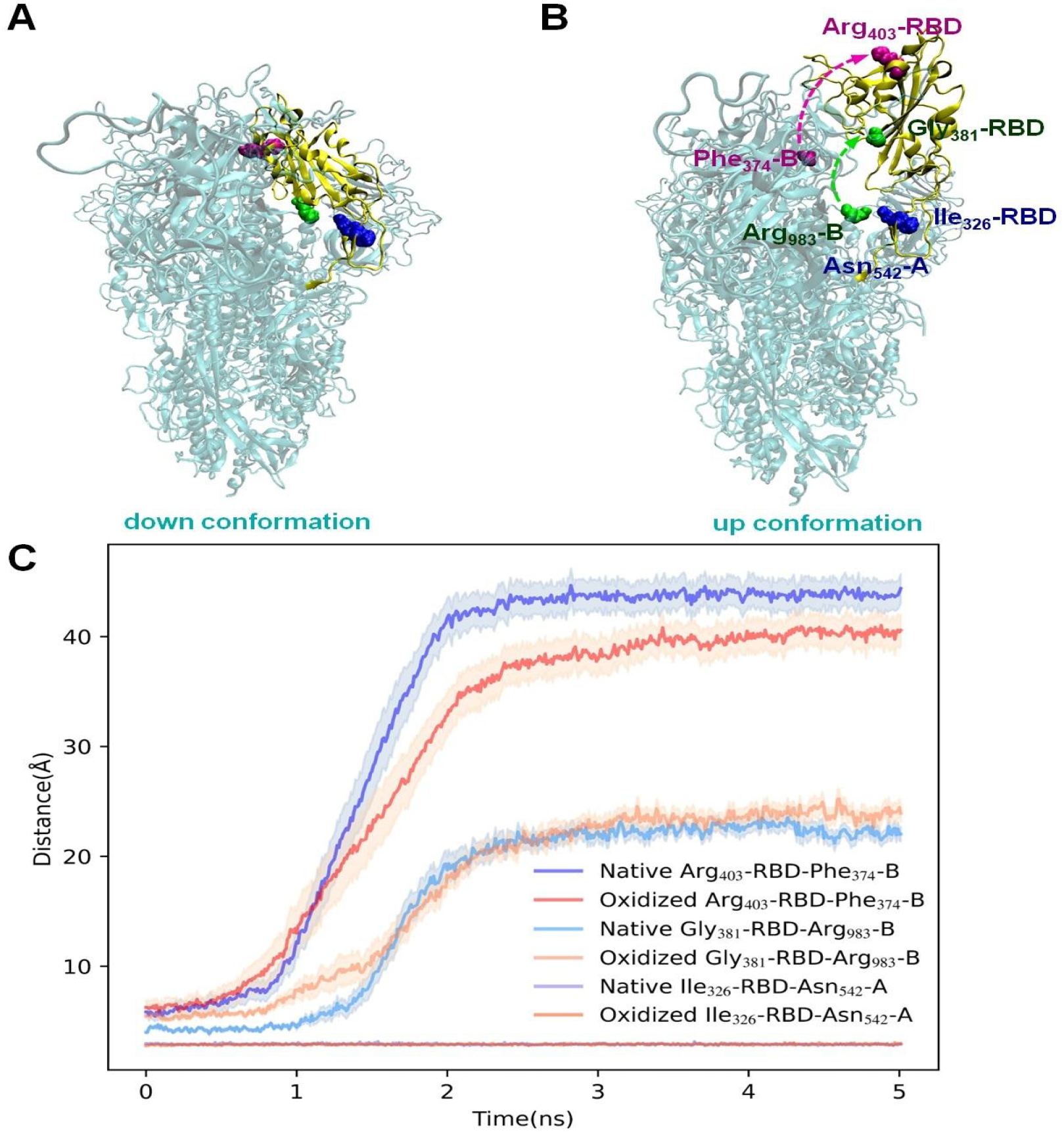
(A) SARS-CoV-2 S trimer in its down conformation, with the RBD domain highlighted in yellow. Three pairs of amino acids forming H-bonds are illustrated: Arg_403_-RBD-Phe_374_-B in magenta, Gly_381_-RBD-Arg_983_-B in green, and Ile_326_-RBD-Asn_542_-A in blue, represented with VDW spheres to emphasize their interactions. (B) Transition to the up conformation of the S trimer, depicting the same amino acid pairs, with arrows indicating the direction of separation. The Ile_326_-RBD-Asn_542_-A pair (blue), located in the hinge domain, remains intact, highlighting its stability. (C) The donor-acceptor distances for these three pairs between the native and oxidized systems throughout the transition. The plot illustrates a less steep slope for the oxidized system, indicating a slower rate of distance change and suggesting fewer dynamic alterations in response to oxidation.

Taken together, these results demonstrate that Cys oxidation renders the conformational transition from the down to up and from the up to down states more challenging, effectively blocking the SARS-CoV-2 S trimer in each respective conformation.

## Summary and Conclusions

Our investigation into the structural dynamics and conformational changes of the SARS-CoV-2 S protein has provided critical insights into its stability and behavior during host cell engagement. Specifically, we focused on the transitions between its down and up states and how these transitions are influenced by Cys residue oxidation. These conformational changes are crucial for viral attachment to host cells, making them a key target for therapeutic intervention.

Using SASA analysis, we identified key residues susceptible to oxidation in the presence of ROS (e.g., generated by CAP treatment). We examined how these modifications affect the SARS-CoV-2 S protein’s conformational transitions through TMD simulations. Our results revealed a significant increase in the energy required for these transitions in the presence of Cys oxidation. Specifically, the work needed for transitioning from the down to the up conformation and vice versa was markedly higher in the oxidized system, i.e., approximately 131 ± 1 kJ.mol^−1^ for the down-to-up transition and 223 ± 6 kJ.mol^−1^ for the up-to-down transition as compared to the native system. This suggests that oxidation stabilizes the SARS-CoV-2 S protein in its current state, either hampering viral entry in the down conformation or increasing susceptibility to neutralization in the up conformation. Consequently, blocking the SARS-CoV-2 S protein in the down state could potentially reduce viral infection, while blocking it in the up-conformation could enhance accessibility to antibodies for neutralization.

The investigation highlights the critical role of glycans in the slower transition and higher work values seen in the oxidized RBD during TMD simulations. The main difference lies in oxidized Cys residues interacting with glycans, which hinders RBD movement and prevents complete upward transition. These interactions contribute to the observed changes in RMSD along the TMD simulations.

Additionally, our analysis of H-bonds and salt bridges before and after oxidation shed light on how these changes contribute to stabilizing the SARS-CoV-2 S protein’s structure in each state. While the overall number of H-bonds remained relatively unchanged post-oxidation, we observed an increase in salt bridge formation, further enhancing the stabilization of the protein’s conformation.

This research advances our understanding of the SARS-CoV-2 S protein’s structural dynamics and underscores the potential of targeting post-translational modifications like oxidation for developing antiviral strategies. By elucidating the energy landscape of conformational transitions and identifying key residues affected by oxidation, our findings provide valuable insights for designing therapeutic interventions aimed at combating SARS-CoV-2 infection and transmission.

## Supporting Information and Data Availability

The Supporting Information and the input files, parameters files and topology files as well as all figures and tables are available free of charge at https://github.com/maryamghasemitarei/Transion-of-SARS-CoV-2-Spike-protein

## Acknowledgments

We sincerely thanks Mohammad Reza Ejtehadi for his constructive feedback and expert guidance throughout the development of this paper.

T.A-N. and M. G. acknowledge support under the European Union – NextGenerationEU Instrument by the Academy of Finland grant 353298.

M. G. gratefully acknowledges the hospitality and support provided by the Abdus Salam International Center for Theoretical Physics (ICTP) (Trieste) during a summer visit.

The computational work was carried out using the Turing HPC infrastructure at the CalcUA core facility of the Universiteit Antwerpen (UA), a division of the Flemish Supercomputer Center VSC, funded by the Hercules Foundation, the Flemish Government (department EWI) and the UA.

